# Crosstalk between the Ino80 complex and TOR signaling drives fungal adaptation to hypoxia through chromatin remodeling

**DOI:** 10.64898/2025.12.19.695452

**Authors:** Manjari Shrivastava, Faïza Tebbji, Morgane Dubé, Antony T. Vincent, Adnane Sellam

## Abstract

In the human host, the opportunistic yeast *Candida albicans* must adapts to niches that vary markedly in oxygen tension, ranging from the hypoxic gastrointestinal and vaginal tracts to more oxygen-rich niches such as the skin. Thriving across this spectrum requires the capacity to dynamically rewire core metabolic pathways in response to fluctuating oxygen levels. Yet, despite the central role of oxygen in shaping fungal metabolism and fitness, the genetic basis underlying hypoxic adaptation in *C. albicans* remains incompletely understood. Here, we performed a genetic screen to identify genes required for adaptation to hypoxia (5% O_2_). This comprehensive approach revealed numerous genes spanning diverse metabolic and stress-related functions that are essential for optimal growth under oxygen-limiting conditions. Notably, this study uncovers a previously unrecognized role for the TOR (Target of Rapamycin) signaling pathway and the chromatin-remodeling complex Ino80 in orchestrating the fungal adaptive response to hypoxia. Integration of *ino80* mutant transcriptomic data with Ino80 DNA-binding and nucleosome-occupancy profiles under conditional *INO80* depletion revealed that this chromatin remodeler regulates the cellular phosphate demand associated with hypoxic exposure. The similarity between chromatin accessibility changes following *TOR1* and *INO80* depletion further suggests that Ino80 acts in concert with the TOR pathway to support growth and maintain phosphate homeostasis under low-oxygen conditions. Collectively, these findings provide new insight into how nutrient and oxygen signaling converge on chromatin remodeling, offering a mechanistic framework for understanding the regulatory logic underlying hypoxic adaptation in pathogenic fungi.

**Importance:** *Candida albicans* is an opportunistic fungal pathogen that colonizes diverse host niches, many of which are oxygen-limited. Understanding how this organism adapts to hypoxia is critical for elucidating the mechanisms underpinning both commensalism and pathogenicity of this yeast. Here, we reveal that the TOR signaling pathway and the Ino80 chromatin-remodeling complex are key regulators of fungal growth under low-oxygen conditions. Our data show that Ino80 coordinates phosphate homeostasis during hypoxia and functions in concert with TOR to maintain chromatin accessibility and metabolic adaptation. Our results uncover how oxygen and nutrient cues converge on chromatin remodeling, providing a framework for understanding how *C. albicans* and other human-associated fungi modulate their physiology to thrive and evolve in diverse host environments.

## Introduction

Molecular oxygen (O_2_) underpins cellular life, serving as an essential substrate for key metabolic reactions and sustaining cellular bioenergetics. As the terminal electron acceptor in mitochondrial respiration, O_2_ drives ATP generation and supports a multitude of O_2_-dependent enzymatic reactions, including lipid desaturation, oxidative protein folding, sterol and porphyrin (heme) biosynthesis (1–4). Oxygen is also essential for multiple metabolic reactions leading to the synthesis of nicotinamide adenine dinucleotide (NAD), an important co-factor for many enzymes and a critical regulator of cellular redox homeostasis (5). Human-associated fungi such as the opportunistic yeast *Candida albicans* occupy different niches with contrasting abundance of O_2_. Except for the skin and other superficial normoxic sites, most anatomical environments encountered by *C. albicans* are characterized by low O_2_ tension, ranging from approximately 1-5 % O_2_ in mucosal surfaces such as the gastrointestinal and vaginal tracts to near-anoxic conditions within deep tissues and biofilm microenvironments (6–8). Therefore, to maintain fitness within the host, *C. albicans* must dynamically readjust its metabolism to limit dependence on O_2_-requiring processes and preserve energy and redox equilibrium under O_2_-limiting conditions (9).

Adaptation to fluctuating O_2_ availability requires not only metabolic flexibility but also precise regulatory systems that can perceive and transduce hypoxic signals into coordinated transcriptional and physiological responses. A prevailing model posits that O_2_ sensing in fungi relies on intracellular levels of O_2_-dependent metabolites, including ergosterol, unsaturated fatty acids, and heme (10, 11). Yet, recent metabolomic analyses suggest that this model may be oversimplified, as depletion of O_2_-dependent lipids for instance was evident only at later stages of hypoxic exposure, rather than during the initial response (9). Thus, early hypoxia perception may not rely primarily on O_2_-dependent metabolites. In *S. cerevisiae*, heme availability functions as a key O_2_ sensor (12). Under aerobic conditions, heme activates the transcription factor Hap1, which promotes respiration gene expression while inducing the repressors Rox1 and Mot3 to silence hypoxia-responsive genes. Conversely, under low O_2_ conditions, the decline in heme levels inactivates Hap1, releasing Rox1- and Mot3-mediated repression and thereby enabling hypoxic gene expression. Interestingly, this regulatory architecture is not conserved in all fungi; in *C. albicans* and *Kluyveromyces lactis*, Rox1 does not control hypoxia-associated genes, suggesting rewiring of hypoxia-responsive pathway (13).

While in higher eukaryotes hypoxia is primarily perceived through O_2_-dependent prolyl hydroxylation of HIF (Hypoxia-inducible factor) transcription factors, fungi appear to employ similar strategies (11, 14). In these organisms, a HIF-analogous regulatory pathway relies on the sterol regulatory element-binding protein (SREBP) transcription factor, which plays a central role in adaptation to hypoxia (11, 15). In *Schizosaccharomyces pombe*, the prolyl hydroxylase Ofd1 regulates SREBP (Sre1) stability, promoting its degradation in the presence of O_2_ and allowing its accumulation under hypoxia (15, 16). Additionally, in response to sterol or O_2_ depletion, Sre1 undergoes proteolytic cleavage that releases it from Golgi to enter the nucleus (11). In the human pathogenic fungi *Aspergillus fumigatus, Histoplasma capsulatum* and *Cryptococcus neoformans*, SREBPs perform similar roles in hypoxic adaptation and are essential for fungal virulence (17–19). In *C. albicans*, no functional SREBP homolog has been identified; instead, the zinc-finger transcription factor Upc2 mediates a comparable function by regulating ergosterol biosynthesis in response to O_2_ limitation (20, 21). However, genetic inactivation of *UPC2* in *C. albicans* results in only a mild phenotype under hypoxic conditions, in contrast to mutants in *S. pombe*, *A. fumigatus*, *H. capsulatum* and *C. neoformans*, where loss of SREBP function leads to severe growth defects or lethality under O_2_ limitation (17–19, 22). These observations highlight that, while SREBP-dependent transcriptional programs are critical for hypoxic adaptation in many fungi, the upstream mechanisms that sense O_2_ limitation and orchestrate metabolic reprogramming in SREBP-lacking species, such as *C. albicans* and several related ascomycetes of medical and industrial relevance, remain largely undefined.

We previously screened a targeted mutant library of *C. albicans* transcriptional regulators and identified Snf5, a core subunit of the SWI/SNF chromatin-remodeling complex, as a critical determinant of fungal fitness under hypoxic conditions (23). In the present study, we expanded this approach by leveraging the GRACE (Gene Replacement and Conditional Expression) collection, which comprises 2,368 conditional mutants representing approximately 40% of all predicted *C. albican*s genes (24). Through this comprehensive screen, we identified 48 genes spanning diverse functional categories that are required for optimal growth under hypoxia, revealing new pathways likely involved in O_2_ sensing, signaling, and metabolic adaptation to O_2_ depletion. Overall, this study reveals a previously unrecognized role of the TOR (Target Of Rapamycin) signaling pathway in controlling the adaptive response of *C. albicans* to hypoxia. We further identified two subunits of the INO80 chromatin-remodeling complex as critical for sustaining fungal growth under O_2_-depleted conditions. Integration of *ino80* mutant transcriptomic data with INO80 DNA-binding and nucleosome-occupancy profiles under conditional *INO80* depletion revealed that this chromatin remodeler controls the cellular phosphate demand accompanying hypoxic exposure. The similarity between chromatin accessibility changes upon *TOR1* and *INO80* depletion suggests that Ino80 functions in concert with the TOR pathway to promote growth and maintain phosphate homeostasis under O_2_-limiting conditions. These findings provide new insight into how nutrient and O_2_ signaling converge on chromatin remodeling mechanisms, offering a foundation for understanding the regulatory logic that underlies hypoxic adaptation in pathogenic fungi.

## Results

### Genetic survey for fungal cell fitness modulators under hypoxic environment

To identify genes required for fungal fitness under O_2_-depleted environment, we screened the GRACE (24) collection of 2368 conditional mutants in *C. albicans*. Cells of the GRACE collection, where an ORF was deleted at one locus and the 5’cis-regulatory region of the other locus was replaced by a tetracycline-repressible promoter, were grown under both normoxic (21% O_2_) and hypoxic conditions (5 % O_2_) for 24 hours in the presence of doxycycline (**Figure 1A**). To identify mutants with a specific growth defect under hypoxia, we calculated the differential growth score (DG) measuring the difference between the relative growth of each mutant to the WT under hypoxia and normoxia (DG= [OD ratio (mut/WT) _hypoxia_ - OD ratio (mut/WT) _normoxia_] x100). Mutants with a growth pattern similar to that of the WT strain or exhibiting the same defects in both hypoxia and normoxia have DG scores of ∼ 0. Empirical DG cutoffs of ± 9 were applied to identify mutants specifically impaired (DG ≥- 9) or buffered (DG ≤ +9) under hypoxia. This empirical cutoff values were determined based on a benchmark phenotype of *cyr1*, a mutant of the adenylyl cyclase, which was previously shown to exhibit an exacerbated growth impairment under hypoxia as compared to normoxic conditions (23). Using this criterion, we identified 48 mutants with exacerbated growth under low oxygen levels and five whose growth was rescued by hypoxia (**Figure 1B** and **Supplementary Table S1**). Pathway-enrichment analysis uncovered that genes required for hypoxic growth were predominantly enriched in vesicle-mediated trafficking and processes related to metabolism including carbon utilization, lipid, and amino acids biosynthesis (**Figure 1C)**. As expected, several oxygen-dependent metabolic genes, such as those involved in ergosterol (*ERG11*, *ERG8*) and heme biosynthesis (*HEM2*, *HAP31*), were among the important hypoxic determinants (**Supplementary Table S2** and **Figure S1**). Interestingly, a significant subset of hypoxia-sensitive mutants carried disruptions in genes encoding chromatin-associated factors and DNA-binding proteins (**Figure 1C-D**). These included *SWI1*, a component of the SWI/SNF chromatin-remodeling complex previously shown to modulate metabolic adaptation and antifungal susceptibility under hypoxia (23, 25), as well as *INO80* and *IES1*, encoding subunits of the INO80 complex (**Figure 1D**). The screen also identified factors involved in amino acid biosynthesis, nutrient sensing (e.g., *KOG1*, a component of the TORC1 complex) (26), cell wall organization, DNA repair, and ribosome biogenesis as key contributors to *C. albicans* growth under hypoxic conditions (**Supplementary Table S2**). Interestingly, hypoxia mitigated the growth defects of five mutants, three of them (*cbf1*, *ras1*, and *slm2*), also displayed a severe growth defect under normoxia (**Figure 1B**). These genes encode the transcription factor Cbf1, which regulates ribosomal biogenesis, the RAS-associated GTPase Ras1, and Slm2, a conserved component of the TORC2 complex in *S. cerevisiae*. Collectively, this screen demonstrates that hypoxic growth depends on a broad network of metabolic and regulatory pathways which likely reflect the complex physiological reprogramming required for *C. albicans* survival in O_2_-limited host niches.

**Figure 1.**
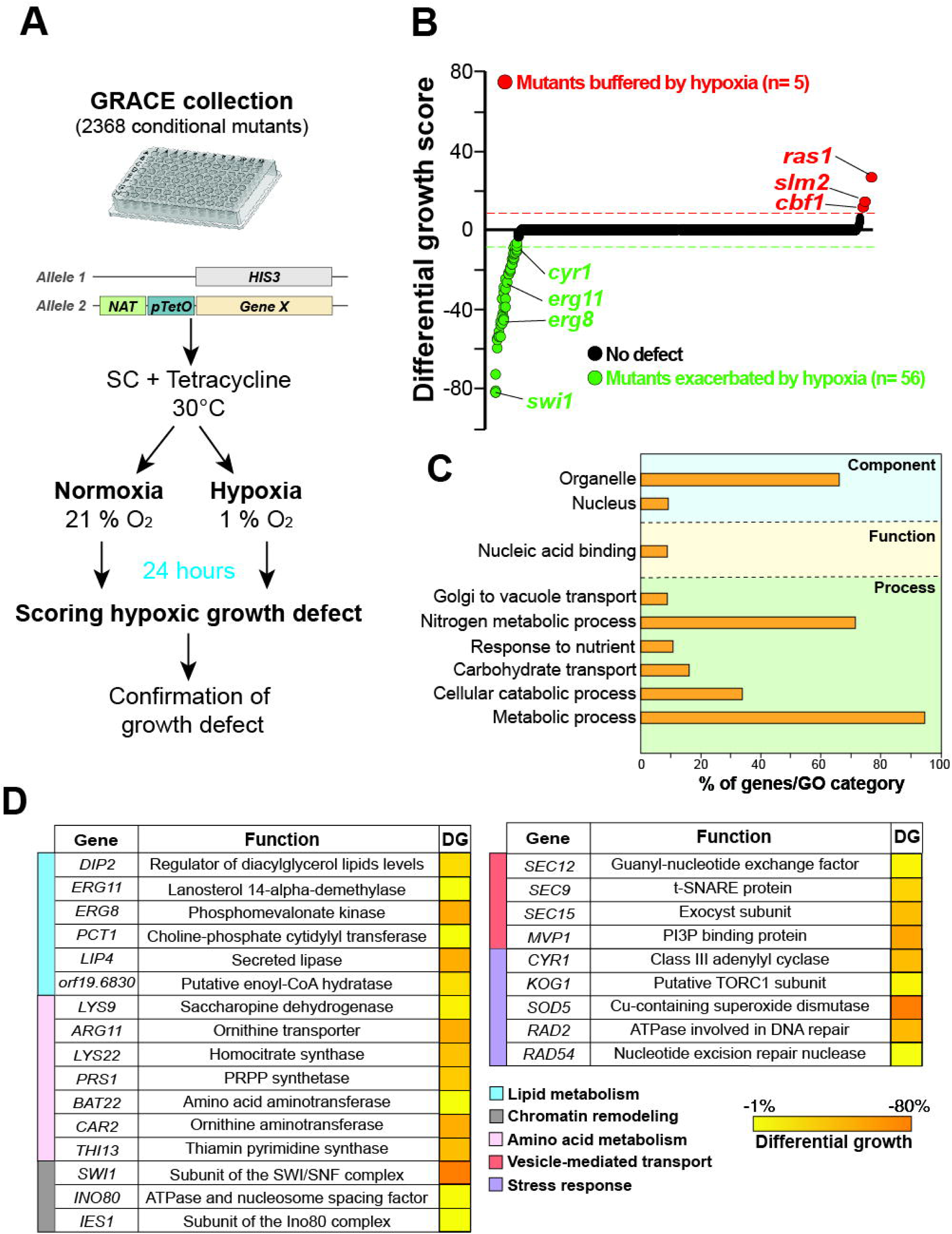
Genetic requirements for *C. albicans* growth under hypoxic conditions. (**A**) Experimental workflow of the genetic screen. Growth (OD_600nm_) of individual mutants of the *C. albicans* GRACE collection was assessed in SC medium at 30°C in both normoxic (21% O_2_) and hypoxic (1% O_2_) conditions. (**B**) Mutants ranked by their growth defect under hypoxia. Differential growth (DG= [OD ratio (mut/WT)_hypoxia_ - OD ratio (mut/WT)_normoxia_] × 100) was assessed for each mutant. Red and green dots represent mutants buffered (DG > −9) and exacerbated (DG < 9) by hypoxia, respectively. (**C**) GO term enrichment of mutants exhibiting growth defects under hypoxia. (**D**) Subset of enriched biological functions of the 48 genes required for fungal hypoxic growth.

### Modulation of TOR activity is required for hypoxic adaptation

We found that the *kog1* GRACE mutant exhibited a significant growth defect under hypoxia as compared to normoxic conditions suggesting a potential role of the TOR pathway in the modulation of fungal growth under O_2_-depelted environments (**Figure 1D**). Consistent with this, progressive reduction of *KOG1* expression led to a dose-dependent impairment of growth specifically under hypoxia as doxycycline concentrations increased (**Figure 2A**). Additionally, all genes of the *C. albicans* TOR complex (TORC1) were haploinsufficient under hypoxia (**Figure 2B**). Furthermore, growth reduction by the inhibitor of the TOR pathway, rapamycin, was exacerbated under hypoxia as compared to normoxic conditions (**Figure 2C)**. Of note, transition of *C. albicans* WT cells from normoxia to hypoxia is accompanied by a moderate growth reduction (**Figure 2C-D**). This growth attenuation was absent in the *TOR1-1* strain, which carries a mutation conferring rapamycin resistance and impairing TOR-mediated nutrient sensing (27) (**Figure 2D**). In addition to exhibiting a proliferative advantage under O_2_-depleted conditions, the *TOR1-1* mutant failed to display the rapamycin hypersensitivity observed in the WT under hypoxia (**Figure 2E**). To further assess TOR activity, we monitored the phosphorylation status of the ribosomal protein S6 (Rps6), a conserved downstream distal effector of the TOR pathway (28, 29). Rps6 phosphorylation decreased under hypoxic conditions, indicating that TOR signaling is repressed in response to low O_2_ availability (**Figure 2F**). Gene set enrichment analysis (GSEA) of transcriptional profiles from hypoxia-exposed *C. albicans* cells (30) revealed a significant overlap with expression signatures characteristic of reduced TOR activity (31) (**Figure 2G**). Collectively, these findings demonstrate that reducing TOR signaling is a critical component of *C. albicans* adaptation and growth under hypoxic conditions.

**Figure 2.**
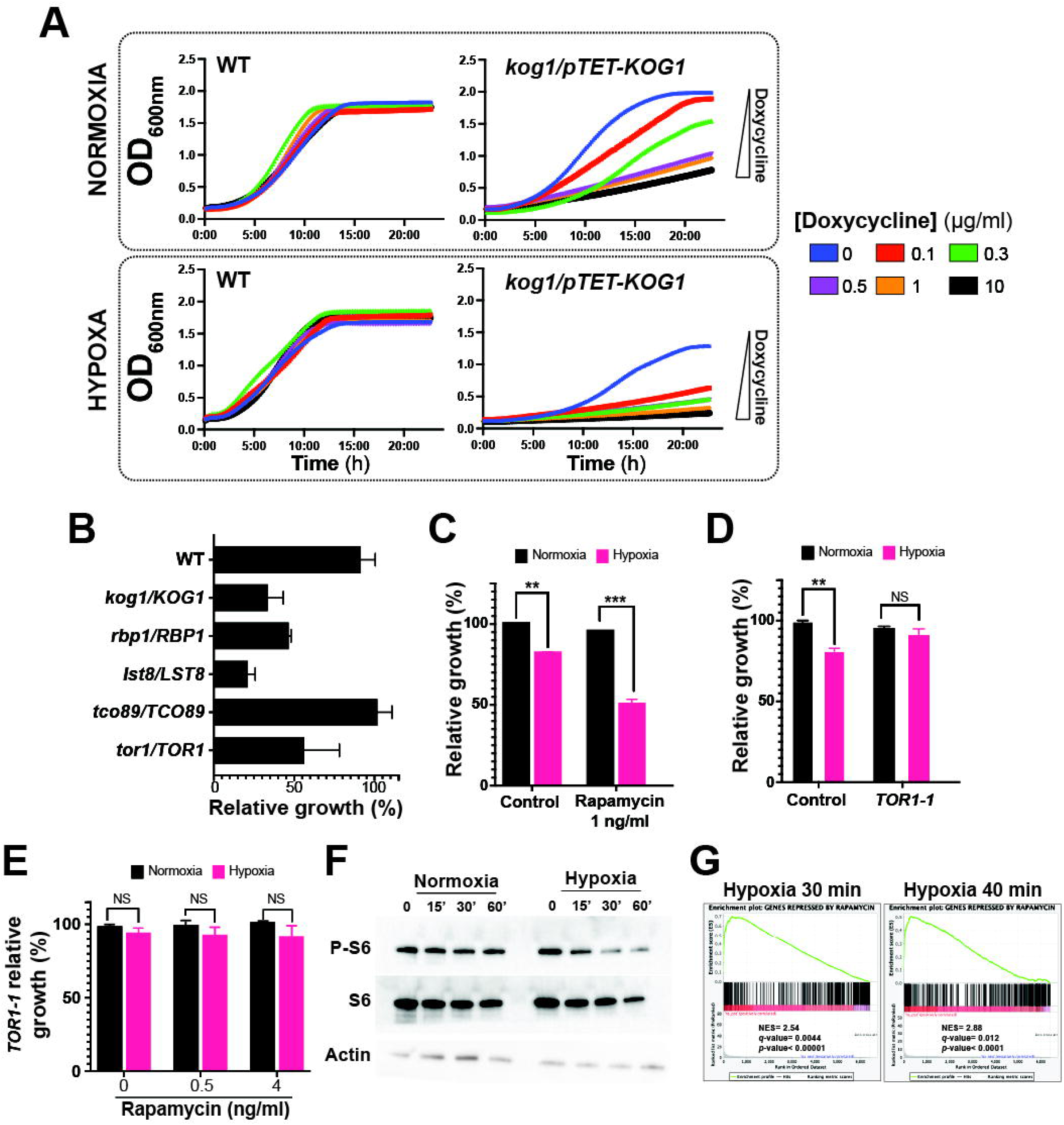
TOR activity modulates fungal growth under hypoxia. (**A**) Overnight cultures of the WT (CAI4) strain and cells expressing *KOG1* from the *tetO* promoter (*kog1*/pTET-*KOG1*) were pre-grown in SC medium and inoculated in SC medium containing increasing concentrations of doxycycline (0.1, 0.3, 0.5, 1 and 10 µg/ml) under both hypoxic and normoxic conditions, with OD_600_ readings taken every 10 minutes. (**B**) Reduced dosage of *C. albicans* TOR pathway genes causes growth defects under hypoxia. Heterozygous mutants (*tor1*/*TOR1*, *kog1*/*KOG1*, *rbp1*/*RBP1*, *lst8*/*LST8*, *tco89*/*TCO89*) and the WT strain (CAI4) were grown in SC medium at 30°C under both normoxic (21% oxygen) and hypoxic conditions (5% oxygen) for 16 h. (**C**) Effect of hypoxia and rapamycin treatment on the *C. albicans* SC5314 strain. Cells were grown for 24 hours under either hypoxia or normoxia in the presence or absence of 1 ng/ml of rapamycin. (**D-E**) Impact of hypoxia (**D**) and TOR inhibition (**E**) on the rapamycin-resistant mutant *TOR1-1*. WT (SC5314) and *TOR1-1* strains were grown for 24 hours under either hypoxia or normoxia in the presence or absence of rapamycin. Cells were treated as described in (**B**). The hypoxic relative growth in (**B**-**E**) was calculated as the OD ratio of the hypoxic to the normoxic cultures and is expressed as a percentage. The results represent the means of the results from at least three biological replicates. Statistical significance was tested using Student’s t-test and is indicated as follows: **, P < 0.01; ***, P < 0.001. NS, not significant. (**F**) Western blot analysis of Rps6 phosphorylation (P-S6) under hypoxia. *C. albicans* SC5314 cells were exposed to hypoxia or normoxia for the indicated times. Cell lysates were immunoblotted for P-S6 and total protein levels of Rps6 and actin (loading control). (**G**) GSEA analysis revealed a significant correlation between cells exposed to hypoxia for 30 and 40 minutes and cells treated with rapamycin. Each bar corresponds to one gene. NES, normalized enrichment score.

### Global chromatin dynamic is altered by Ino80 loss under hypoxic environment

We previously uncovered that the chromatin remodeling complex SWI/SNF couples carbon metabolic flexibility with growth under hypoxia (23). Consistent with this, our genetic screen identified a mutant of the SWI/SNF subunit Swi1, which displayed a pronounced growth defect under hypoxic conditions (DG = 82%). In addition, mutants of two subunits of the chromatin remodeling complex INO80 (*ino80* and *ies1*) were also uncovered, underscoring the importance of chromatin regulation in controlling growth under hypoxia (**Figure 1D**). To investigate the role of Ino80 in hypoxic adaptation, we first generated an *ino80* deletion mutant and confirmed its growth defect under hypoxia, similar to that of the corresponding conditional mutant (**Figure 3A**). We then analyzed the *ino80* transcriptome by RNA-seq and compared it with the WT under the same conditions. Downregulated transcripts in *ino80* were predominantly enriched in processes related to ion transport, amino acid catabolism and cell adhesion (**Figure 3B** and **Supplementary Table S3**). In contrast, genes involved in sulfur metabolism, biotin biosynthesis, biofilm formation, and carbohydrate transport were significantly upregulated. To delineate the direct targets of Ino80, we performed ChEC-seq under conditions similar to those used for the RNA-seq analysis. Ino80 occupancy was detected at the promoter regions of eleven downregulated transcripts associated with ion transport and cell wall remodelling (**Figure 3C** and **Supplementary Table S4**). In addition, Ino80 bound to the promoters of sixteen upregulated genes related mainly to biofilm formation and amino acid catabolism (**Figure 3C**). These findings suggest that Ino80 directly mediates both transcriptional activation and repression of distinct biological processes under hypoxia.

**Figure 3.**
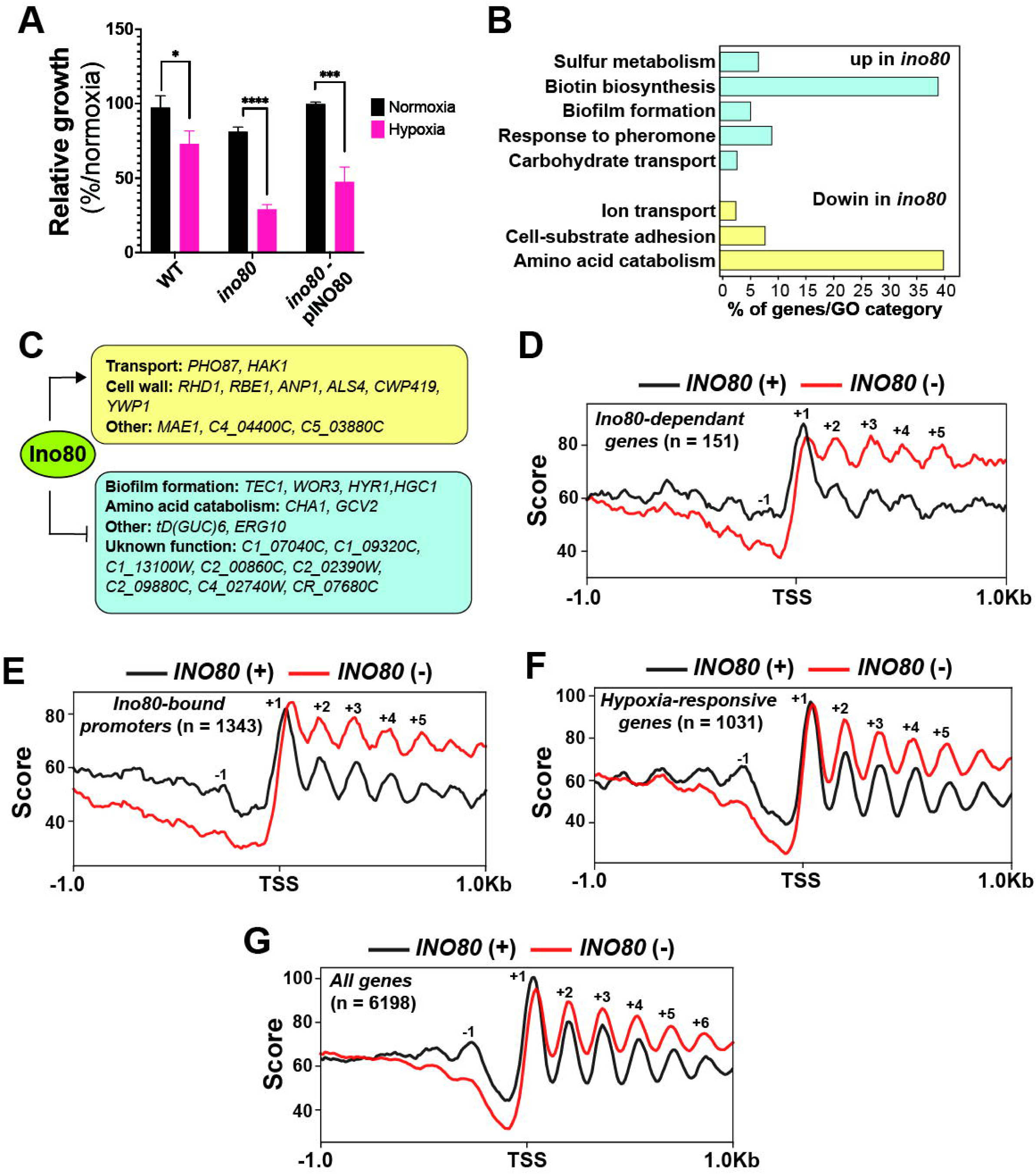
Ino80 regulates nucleosome organization and chromatin accessibility under hypoxic conditions. (**A**) *INO80* inactivation led to a growth defect under hypoxia. Overnight cultures of the WT strain (SN87), the *ino80* mutant and the *ino80* strain complemented with wild type *INO80* (*ino80*-p*INO80*) were grown for 24 hours under either hypoxia or normoxia. (**B**) Gene functions and biological processes enriched in the transcriptional profiles of the *ino80* mutant under hypoxia. Differentially modulated transcripts were identified by comparing the *ino80* transcriptome with that of the WT strain (SN87). (**C**) Ino80 hypoxic regulon. An integrated analysis of RNA-seq and ChEC-seq data identifying transcripts that are differentially expressed in the *ino80* mutant and whose promoters are bound by Ino80. Genes upregulated (in blue) or downregulated (in yellow) in the *ino80* under hypoxia are shown, highlighting direct transcriptional targets of the Ino80 complex involved in the hypoxic response. (**D-G**) Diagram illustrating the canonical nucleosomal organization at ±1Kb regions upstream or downstream of the TSS of different sets of genes: gene differentially expressed in the *ino80* under hypoxia (**D**), Ino80-occupied promoters (**E**), *C. albicans* hypoxia-responsive genes (**F**) and all *C. albicans* genes (**G**) in both WT (*INO80* (+)) and *INO80*-depleted cells (*INO80* (−)). Nucleosome positions and their corresponding nucMACC scores were determined using the nucMACC pipeline.

Furthermore, to assess Ino80 contribution on chromatin remodeling at the genome scale under hypoxia, we generated a high-resolution nucleosome map obtained by MNase-seq in *INO80*-depleted cells, referred as *INO80* (−). Nucleosome occupancy was first assessed for genes differentially modulated in the *ino80* mutant and bound by Ino80 under hypoxic conditions. *INO80* depletion led to elevated nucleosome occupancy across coding regions and a concomitant reduction at the promoter regions (**Figure 3D**). We also noticed that the −1 nucleosome was weakly positioned in *INO80* (−) as compared to the WT strain, in addition to a marked displacement of the +1 nucleosome in the direction opposite to the nucleosome-free regions (NFR), consistent with the proposed “puller” activity of the Ino80 complex (32, 33). At the genome scale or when focusing specifically on Ino80-occupied promoters or hypoxia-responsive genes, defined by comparing the WT transcriptome under hypoxia to that of normoxia (**Supplementary Table S5**), similar alterations of nucleosome occupancy were observed (**Figure 3E-G**). Together, these indicate that Ino80 is essential for maintaining global chromatin accessibility that underlies the transcriptional program driving hypoxic adaptation.

### Ino80 modulates growth under hypoxia by controlling the cellular phosphate demand associated with hypoxic adaptation

We found that Ino80 bound the promoter of the low affinity phosphate transporter Pho87 and was required for its activation under hypoxia (**Figure 3C**). Among the *PHO* regulon previously defined (34, 35) and in addition to *PHO87*, Ino80 occupied the promoter of the high-affinity Pi transporters *PHO84* (**Supplementary Table S4**). *PHO84* transcript was also downregulated in *ino80* (Log2FC of −0.56) but did not meet the fold-change cutoff criteria (Log_2_FC of ±1) (**Supplementary Table S3**). As for Ino80-regulated promoters (genes misregulated in *ino80*, with promoter regions bound by Ino80 and exhibiting altered nucleosome organization upon *INO80* depletion), we found that Ino80 was necessary to remodel nucleosome dynamics at the *PHO87* and *PHO84* promoters (**Figure 3D and 4A-B**). Interestingly, unlike other Ino80-regulated promoters, we observed weak nucleosome occupancy within the predicted NFRs of the *PHO87* and *PHO84* promoters in *INO80*-depeleted cells (**Figures 4A-B**). This feature likely corresponds to the previously described “fragile” nucleosomes, which are inherently unstable structures associated with promoters, poising genes for rapid activation in response to environmental cues (36, 37). Thus, Ino80 is likely required to evict these NFR-associated fragile nucleosomes to facilitate *PHO87* and *PHO84* activation under hypoxic conditions.

**Figure 4.**
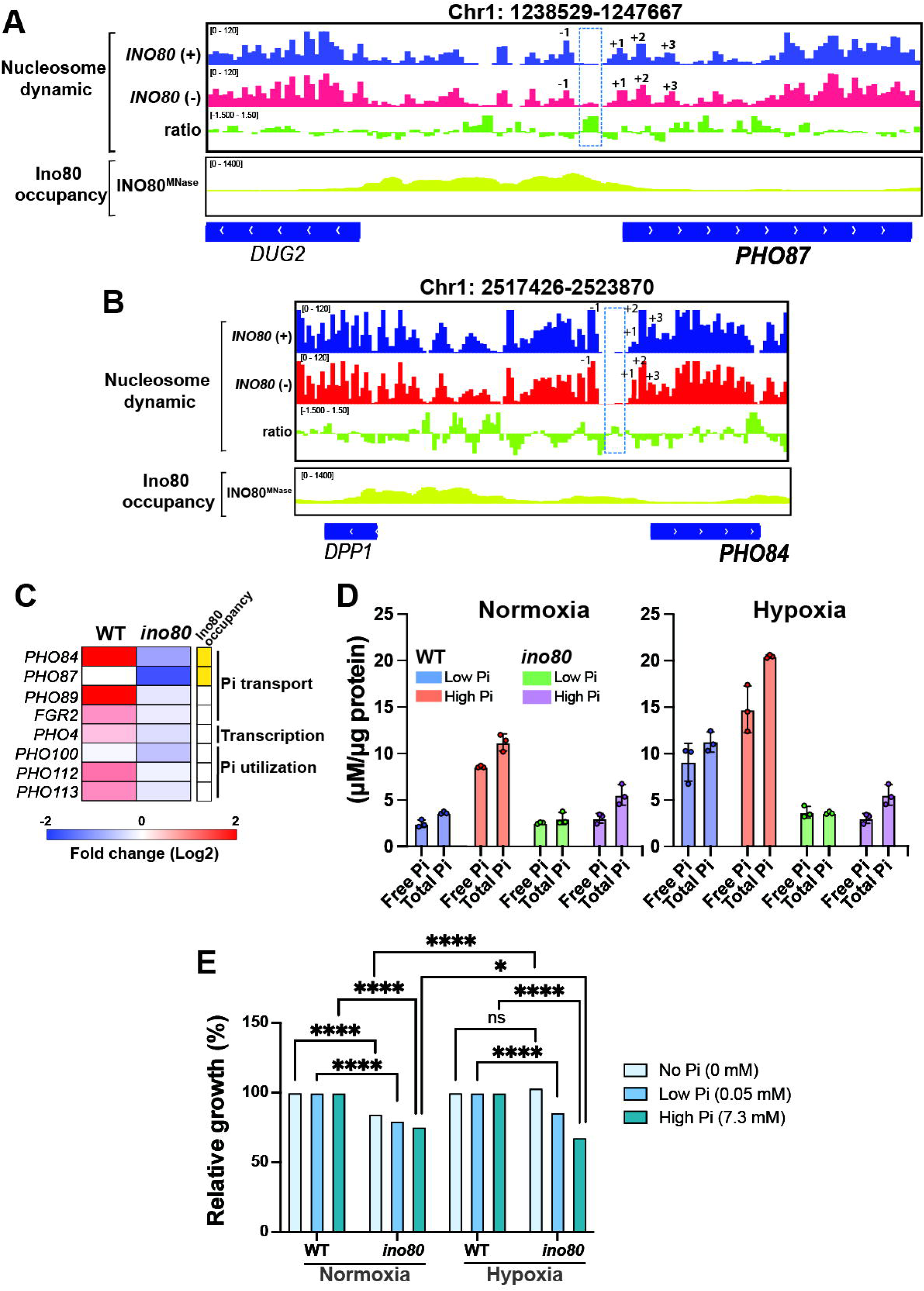
Ino80 modulates hypoxic growth through regulation of phosphate homeostasis. (**A-B**) Nucleosome occupancy maps of *PHO87* (**A**) and *PHO84* (**B**) in WT and *INO80*-depeleted cells. Ino80 binding at the promoter regions, as determined by ChEC-seq is also shown. (**C**) Modulation of the *PHO* regulon by hypoxia in WT strain (SN87) and the *ino80* mutant. Ino80 binding to the promoter of each gene is also shown. (**D**) Free and total inorganic phosphate (Pi) uptake assays in the WT strain (SN87) and the *ino80* mutant grown in SC medium with low (0.05 mM) or high Pi (7.3 mM). (**E**) Relative hypoxic growth of the *ino80* mutant in media containing no, low, or high phosphate (Pi), normalized to the WT (CAI4) growth under the same conditions.

To investigate the contribution of phosphate metabolism to hypoxic adaptation, we first assessed the modulation of the *C. albicans PHO* regulon. In WT cells, transcript of phosphate transporters (*PHO84, 87, 89, FGR2*), phosphatases (*PHO100, 112, 113*), and the transcriptional regulator *PHO4* were upregulated under hypoxia, indicating an increased cellular demand for phosphate during oxygen limitation (**Figure 4C**). In contrast, the *ino80* mutant showed markedly reduced expression of the *PHO* regulon, consistent with an impaired ability to activate phosphate transporters under hypoxic conditions. Direct measurements of intracellular inorganic phosphate (Pi) further supported this defect. In WT cells, hypoxia triggered a substantial increase in both free and total intracellular Pi, regardless of the Pi concentration in the medium (**Figure 4D**). Under normoxia, the *ino80* mutant exhibited reduced intracellular Pi only in high-Pi medium. However, under hypoxia, *ino80* cells accumulated significantly less intracellular Pi than WT, even in low-Pi conditions. Although supplementation with excess phosphate led to a modest increase in Pi levels in *ino80*, these levels remained far below those of WT cells grown under the same conditions (**Figure 4D**). Collectively, these data reveal that Ino80 plays a central role in orchestrating phosphate acquisition to sustain cellular requirements during oxygen limitation.

To further explore the relationship between phosphate availability and hypoxic fitness, we monitored the growth of the *ino80* under hypoxia in media containing no phosphate, low phosphate (0.05 mM), or high phosphate (7.3 mM). Remarkably, the hypoxic growth defect of *ino80* was completely rescued in phosphate-free medium (**Figure 4E**). However, as phosphate concentrations increased, the growth of *ino80* declined in a dose-dependent manner, revealing that phosphate itself becomes detrimental to *ino80* under oxygen limitation. This phosphate-dependent growth pattern contrasts with that of *pho84*, *pho87* and *pho4* mutants, whose growth defects in phosphate-poor medium can be alleviated by high-Pi supplementation under hypoxia (**Supplementary Figure S2).** Together, these findings indicate that Ino80 is required not only for efficient Pi acquisition under hypoxia but also for preventing phosphate-induced toxicity when O_2_ is limiting, highlighting a critical role for Ino80 in maintaining Pi homeostasis during hypoxic adaptation.

### Functional relationship between Ino80 and the TOR pathway in mediating hypoxic adaptation

To test the hypothesis that the TOR pathway contributes to hypoxic adaptation by modulating chromatin dynamics, we mapped the nucleosome landscape of *C. albicans TOR1*-depleted cells (*TOR1* (−)). Overall, at Ino80-occupied loci, *TOR1* depletion led to the disruption of the hypoxic nucleosome accessibility pattern, similarly to *INO80* depletion (**Figure 5A**). A pronounced increase in nucleosome occupancy across coding regions, accompanied by a loss of nucleosome periodicity, particularly at Ino80-regulated promoters, were observed in *TOR1*(−). Moreover, nucleosome occupancy upstream of transcription start sites (TSS) was markedly reduced. Together, these observations highlight the importance of the TOR pathway in linking oxygen status to chromatin remodeling.

**Figure 5.**
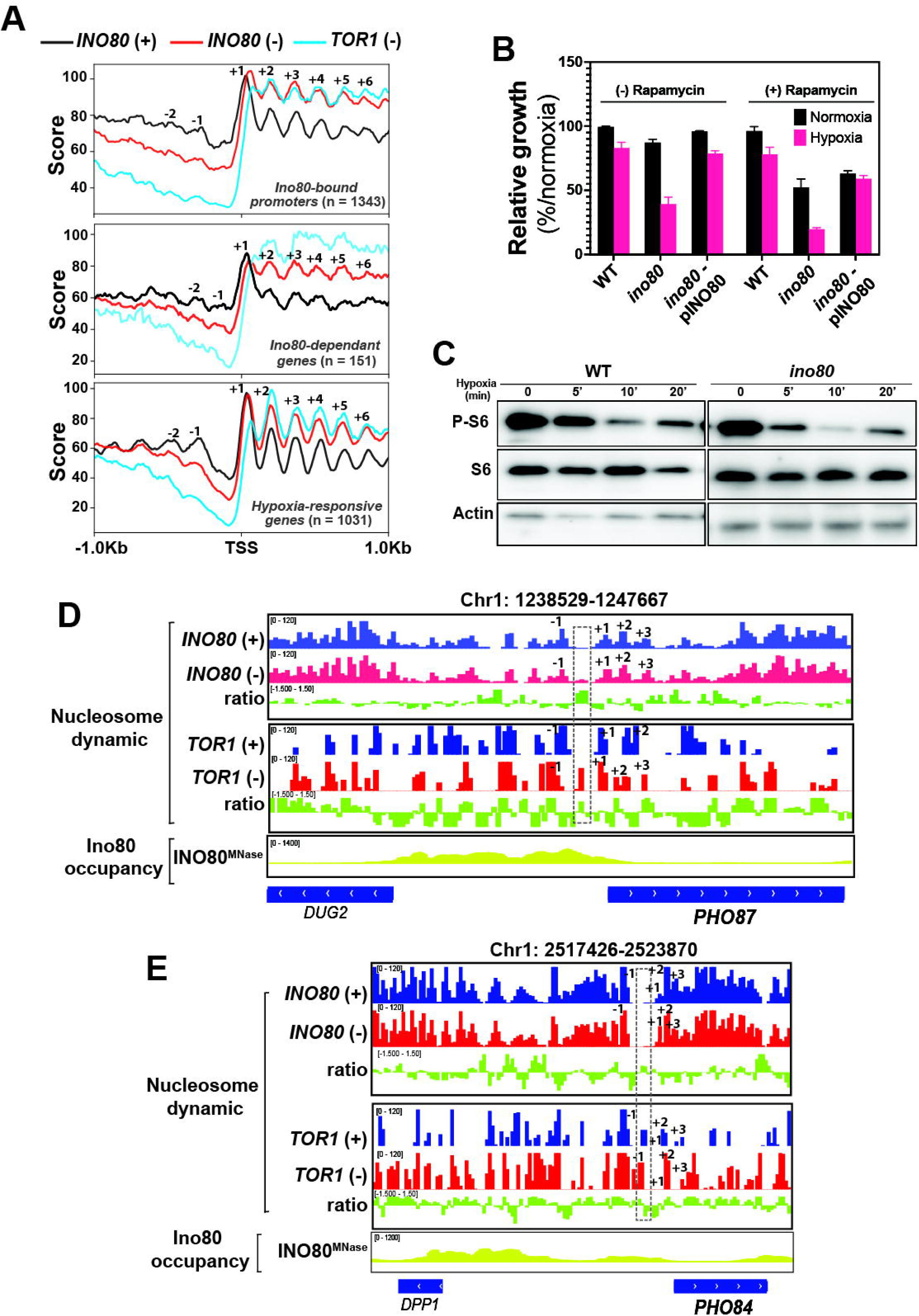
Functional interplay between Ino80 and the TOR pathway in mediating hypoxic adaptation. (**A**) Nucleosome organization at ±1Kb regions upstream or downstream the TSS for different sets of genes, as shown in Figure 3D in WT (*INO80*^+^ and *TOR1^+^*), *TOR1*-depleted and *INO80*-depleted cells (*INO80*^−^ *and TOR1^−^*). Nucleosome position and scores were determined using the nucMACC pipeline. (**B**) TORC1 inhibition exacerbates growth defect of the *ino80* mutant under hypoxia. Overnight cultures of the WT strain (SN87), the *ino80* mutant and the *ino80*-p*INO80* complemented strain were grown for 24 hours under either hypoxia or normoxia in the presence or absence of rapamycin (5 ng/ml). (**C**) Phosphorylation status of the ribosomal protein S6 (Rps6) under hypoxia in the *ino80* mutant. (**D**) Snapshot of nucleosome occupancy and Ino80 binding profiles across the *PHO87* and *PHO84* loci as shown in Figure 4A-B, together with MNase-seq data obtained under *TOR1* depletion. The region indicated by the dashed rectangle highlights the coordinated changes in nucleosome positioning upon *INO80* and *TOR1* depletion.

TOR inhibition produced a nucleosome accessibility profile similar to that observed upon *INO80* depletion, suggesting a functional link between Ino80 and the TOR pathway in regulating the chromatin state under hypoxia (**Figure 5A**). *TOR1* depletion led to a similar nucleosome occupancy pattern as *INO80* (−) with a potential fragile nucleosome at the NFR of *PHO87* and *PHO84* promoters (**Figure 5D-E**). Furthermore, the *ino80* mutant was hypersensitive to rapamycin under both normoxic and hypoxic conditions and exhibited a pronounced reduction in P-S6 phosphorylation under hypoxia, indicating altered TOR activity (**Figure 5B-C**). This indicates that Ino80 acts in concert with the TOR pathway to promote growth under oxygen-limiting conditions.

### Ino80 modulates different *in vivo* fitness attributes

The requirement of Ino80 for fungal virulence was previously demonstrated in a murine model of systemic candidiasis, where the *ino80* mutant was found to be avirulent due to the essential role of this chromatin remodeler in mediating invasive hyphal growth (38). Our current work extends this role by underlying the contribution of Ino80 to adaptation to hypoxic environments, a condition encountered during infection. We confirmed the essential role of Ino80 in fungal virulence, by demonstrating its requirement to infect *Galleria* larvae and to damage human enterocytes (**Figure 6A-B**). We also showed the conserved role of Ino80 in inositol prototrophy, similar to its ortholog in budding yeast (39) (**Figure 6C**). Consistent with the altered transcript levels of genes associated with adhesion and biofilm formation, the *ino80* mutant displayed a pronounced defect in biofilm formation, particularly under hypoxic conditions (**Figure 6D**). Furthermore, *ino80* exhibited a reduced tolerance to increased temperature and was essential for growth at 37°C under hypoxia (**Figure 6E**). Overall, these findings suggest that Ino80 is a pleiotropic factor that regulates multiple biological processes critical for fungal fitness *in vivo*.

**Figure 6.**
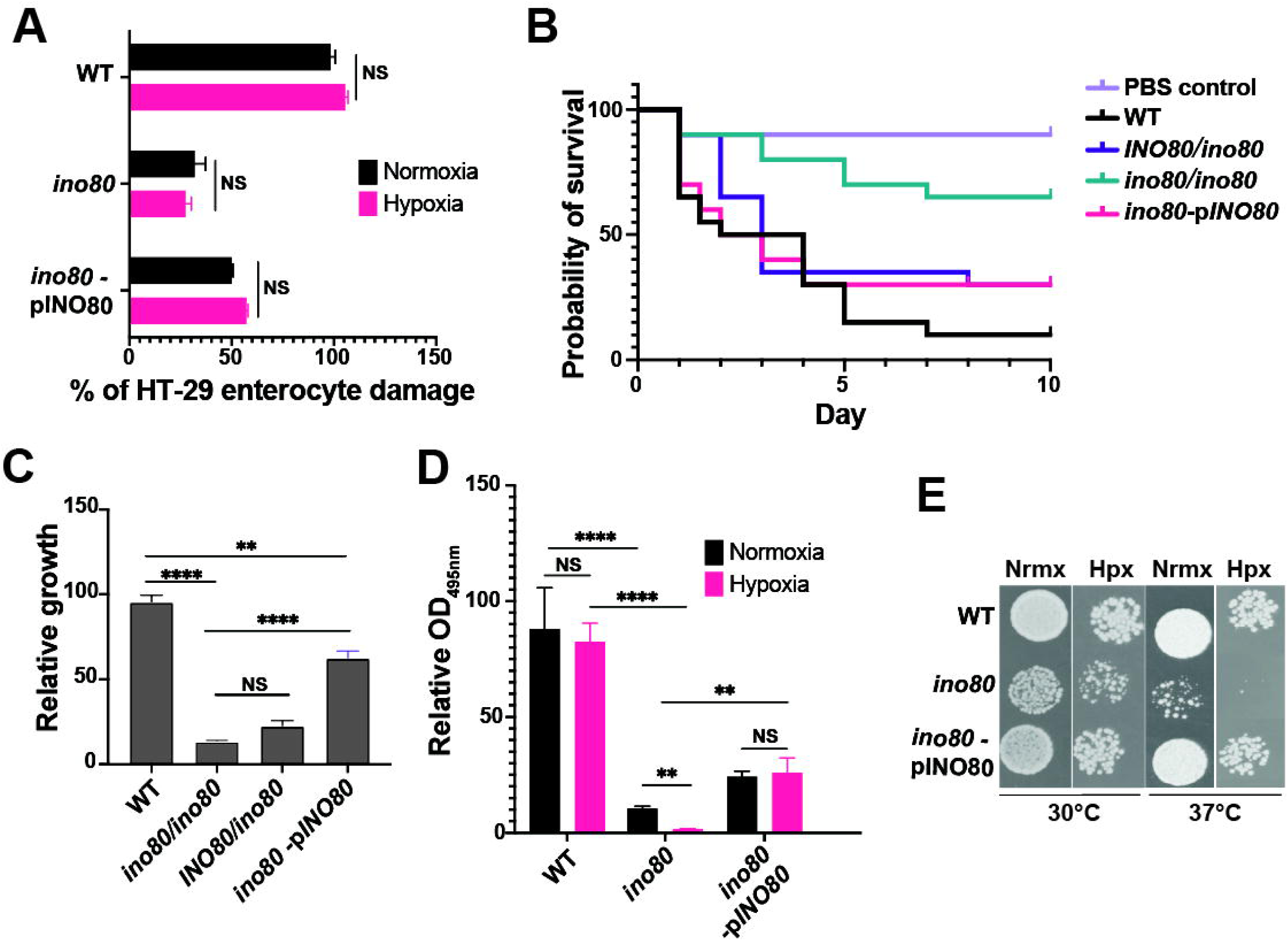
Ino80 modulates different critical function for *in vivo* fitness and virulence. (**A-B**) Ino80 is required for fungal virulence. The ability of the *ino80* mutant, the parental WT (SN87) and the complemented strain (*ino80*-p*INO80*) was tested to damage to human HT-29 enterocytes under both normoxic and hypoxic conditions (**A**). Cell damage was assessed using the lactate dehydrogenase (LDH) release assay and was expressed as a percentage of LDH activity. (**B**) *Galleria* systemic infection assay. WT (SN87), heterozygous (*INO80*/*ino80*) and homozygous (*ino80*/*ino80*) *ino80* mutants, together with the PBS control, were injected into *G. mellonella* larvae, and survival was monitored daily for 10 days. (**C**) *ino80* auxotrophy for inositol. WT (SN87), heterozygous (*INO80*/*ino80*) and homozygous (*ino80*/*ino80*) *ino80* mutants, and the complemented strain were grown in SC medium lacking inositol for 24 hours. (**D**) Ino80 is essential for biofilm formation under hypoxia. *C. albicans* cells were seeded into 96-well polystyrene plates and incubated at 37°C for 3 hours. Biofilm biomass was assessed using the XTT metabolic assay. (**E**) Ino80 is required for tolerate normal body temperature. Spot assays of the *ino80* mutant together with the WT and the complemented strains at 30°C and 37°C under both normoxia and hypoxia.

## Discussion

This study offers the first comprehensive delineation of the genetic determinants that underpin hypoxic adaptation in a fungus. Consistent with previous studies, our analysis identified known regulators of hypoxic metabolic reprogramming and fitness, including the SWI/SNF complex and Cyr1 (23). We also recovered genes predicted to contribute to hypoxic adaptation due to their O_2_-dependent functions in ergosterol and heme biosynthesis. Beyond these findings, we identified many genes involved in other metabolic pathways, including lysine and arginine biosynthesis, lipid and carbohydrate metabolism. These findings demonstrate that hypoxia triggers coordinated adjustments across diverse biosynthetic routes and support previous reports of extensive metabolome remodeling in *C. albicans* (9, 23). Hypoxia is also known to induce substantial remodeling of the *C. albicans* cell wall, most notably through reduced β-glucan exposure, a strategy that promotes evasion of host immune detection (9, 40). Consistent with this, our screen revealed the involvement of two cell-wall glucosyltransferases (*ALG8* and *OCH1*) and a glycosidase (*SUN41*), which may contribute to these O_2_-dependent structural modifications. While the role of these genes in cell wall masking has not been examined in *C. albicans*, studies in *S. cerevisiae* have shown that *OCH1* was required for masking β-glucan and immune recognition (41). Stress-responsive genes such as the superoxide dismutase *SOD5* and DNA replication and repair genes (*CDC45*, *TOF1*, *RAD54*) were also identified. These findings likely reflect, respectively, the need to mitigate oxidative stress and the previously reported rate-limiting impact of nucleotide scarcity under hypoxia (9). Because hypoxia constrains *de novo* nucleotide biosynthesis, cells experience impaired DNA synthesis and increased replication stress, thereby elevating the demand for replication and repair factors to maintain genome integrity (42, 43). Overall, our screen uncovered novel modulators of hypoxic fitness in *C. albicans*, providing new avenues for dissecting the molecular basis of adaptation to low O_2_ levels and metabolic readjustments.

Our study points to the TOR pathway as a key player in hypoxic adaptation in *C. albicans*. The observed reduction in Rps6 phosphorylation, along with a transcriptional profile resembling rapamycin-treated cells, indicates that hypoxia attenuate TOR activity, which may contribute to the broader metabolic remodeling and stress response observed under low O_2_ conditions (9, 30). In budding yeast, diminished TOR activity in response to environmental stress and nutrient deprivation is known to derepress multiple signaling and transcriptional adaptive programs that promote cell survival (44). Thus, a comparable TOR-dependent regulatory cascade may operate in *C. albicans*, modulating metabolic reprogramming and growth to support fitness in O_2_-poor niches. While more than a dozen direct or distal TOR effectors have been characterized in *S. cerevisiae* (45), their counterparts in *C. albicans* remain unknown. Defining these TOR targets will be essential for determining which downstream pathways are derepressed under hypoxia. The possibility that TOR modulates hypoxic adaptation through Ino80 as a downstream effector is supported by several independent lines of evidence. First, the nucleosome alterations detected in *TOR1*-depleted cells, both at the *PHO87* and *PHO84* promoters and across Ino80-dependent loci, mirror the chromatin defects observed in *INO80*-depleted cells, suggesting that TOR activity influences Ino80-mediated promoter remodeling. This similarity implies that Ino80 may execute part of the TOR-dependent control of nucleosome architecture during O_2_depletion. Second, recent studies in *C. albicans* demonstrate that phosphate homeostasis and activation of the *PHO* regulon are regulated by TOR signaling (46). As our data show that Ino80 is required to meet the elevated phosphate demand under hypoxia, this remodeler may function as a key integrator of TOR signaling and *PHO* pathway activation, positioning this remodeling complex as a regulatory hub coordinating phosphate sensing, chromatin structure, and hypoxic transcriptional reprogramming. Furthermore, because increased intracellular phosphate transport in *C. albicans* has been shown to activate and sustain TOR activity as part of a feedback control mechanism (46), the impaired phosphate acquisition observed in the *ino80* mutant may weaken this regulatory loop, thereby diminishing TOR activity under hypoxic conditions. Consistent with this model, our data show that the *ino80* mutant exhibits reduced TOR activity under hypoxia and reinforces the idea that Ino80 not only contributes to TOR-dependent chromatin remodeling but may also influence TOR signaling indirectly through its role in phosphate homeostasis. The TOR pathway is a well-established regulator of growth and overall fitness in *C. albicans*, and recent *in vivo* studies demonstrate that modulation of TOR signaling can drive adaptive evolution that supports both commensal fitness and invasive pathogenicity (47, 48). In this context, our findings raise the possibility that TOR functions as an integrative regulatory node that links changes in O_2_ availability to the adaptive evolutionary responses that emerge during host colonization.

Our work revealed that hypoxia exposure leads to an increased cellular demand for phosphate, a response that is mediated by Ino80 and the *PHO* regulon. Several mechanisms could account for this elevated phosphate requirement. Given that ATP hydrolysis releases phosphate and hypoxia reduces intracellular ATP levels (23), the associated decline in ATP may generate a physiological signature resembling phosphate starvation, thereby inducing phosphate acquisition pathways. The increased phosphate demand under hypoxia is further reflected by the elevated levels of various phosphate-containing metabolites of different pathways including glycolysis (e.g., glucose-1-phosphate, glucose-6-phosphate), pentose phosphate pathway (e.g., 5-phosphoribosyl diphosphate), and glycerophospholipids (e.g., glycerol-3-phosphate and phosphatidylinositol), as previously reported (9).

As Ino80 directly regulates the transcription of Pi transporters, we initially expected the *ino80* mutant to phenocopy *pho*-pathway mutants and display impaired growth under phosphate-deprived conditions (35, 46, 49). Unexpectedly, *ino80* cells grew normally in the absence of phosphate and instead exhibited a phosphate-dependent growth defect, with higher Pi concentrations progressively worsening the phenotype. This suggests that the growth defect of *ino80* under hypoxia does not stem from an inability to grow in Pi-limited environments *per se*, but rather from an impaired capacity to handle the acquisition of external phosphate, a demand that is specifically imposed by O_2_ limitation. This may reflect an inability of *ino80* cells to safely store incoming phosphate as vacuolar polyphosphate (34, 50), leading to toxicity when extracellular phosphate is available. Future studies dissecting how Ino80 contributes to managing phosphate influx and storage during hypoxia will be essential to fully understand the mechanistic basis of this phosphate-dependent growth defect.

Previous studies have established a connection between the TOR pathway and Ino80 in modulating various biological functions in *S. cerevisiae*, suggesting a conserved interplay between nutrient signaling and chromatin remodeling across these yeast species (51–54). However, our study also highlights notable differences, reflecting evolutionary divergence between the two yeasts regarding the TOR-Ino80 regulatory axis. For instance, in *S. cerevisiae*, Ino80 has been proposed to act as an effector of the TOR pathway, modulating TOR-responsive genes primarily associated with translation and ribosome biogenesis (51, 52). In contrast, in *C. albicans* our RNA-seq and ChEC-seq data did not capture this regulation. Furthermore, deletion of *INO80* in *S. cerevisiae* was reported to decrease rapamycin sensitivity and increase TOR activity, as measured by Rps6 phosphorylation (51), which is entirely opposite to our findings in *C. albicans*. A specific feature of *C. albicans* Ino80 is that its inactivation leads to a loss of thermotolerance, suggesting a critical role in enabling the fungus to withstand elevated temperatures. Together, our work suggests that Ino80-mediated chromatin remodeling in *C. albicans* modulates distinct biological functions to facilitate adaptation to the host environment.

## Methods

### Strains and growth condition

The *C. albicans* strains used in this study are listed in **Supplementary Table S6**. All strains were routinely cultured at 30°C in yeast-peptone-dextrose (YPD) medium supplemented with uridine (2% Bacto peptone, 1% yeast extract, 2% dextrose, and 50 µg/ml uridine, with the addition of 2% agar for solid medium) or on synthetic complete medium (SC; 1.7% yeast nitrogen base, 0.5% ammonium sulfate, 2% dextrose, 0.2% amino acids, with 50 μg/ml uridine). The *ino80* mutant was constructed from SN152 strain by replacing the entire ORF with a PCR-disruption cassette amplified from the pFA plasmids (55). Complementation of the *ino80* mutant was achieved by amplifying 1 kb upstream of the *INO80* along with the entire ORF. The amplified DNA was cloned into the Cip10 plasmid (56) and the resulting construct was digested with *StuI* and integrated into the *RPS10* locus using the lithium acetate transformation procedure. The GRACE collection (24) was obtained from the National Research Council of Canada research center (NRC Royalmount, Montreal). For a growth screen under hypoxia, GRACE strains and the WT parental strain CAI4 were grown overnight in SC supplemented with 0.05 µg/ml doxycycline for partial gene transcriptional repression (57). Overnight cultures were diluted to an OD_600_ of 0.05 in 96-well plates with fresh SC medium with doxycycline and incubated at 30°C under normoxic (21% O_2_) or hypoxic (1% O_2_) conditions, with OD_600_ readings taken every 10 min for 24 h using a Cytation 5 plate reader. For each strain, growth was assayed in duplicate, and a DG score was given (DG = [OD ratio (mut/WT) _hypoxia_ - OD ratio (mut/WT)] _normoxia_ × 100). Yeast Nitrogen base without amino acids, without ammonium sulfate, and without Inositol (CYN3801, Formedium), was used to test *ino80* inositol auxotrophy. Phosphate-defined media at acidic pH (pH 3) were prepared using a phosphate-free Yeast Nitrogen Base (CYN6802, Formedium) supplemented with KCl and titrated with the indicated concentrations of KH_2_PO_4_. Spot dilution assays were performed as previously described (25).

### Western blot

Ribosomal protein S6 (Rps6) phosphorylation was assessed as previously described (48). Overnight cultures of *C. albicans* strains were grown in YPD at 30°C with shaking, washed with PBS, and diluted into fresh SC medium to an OD_600_ of 0.2. After 2 h, cells were either subjected to hypoxic conditions or treated with 5 ng/ml rapamycin for the indicated durations. Cells were harvested by centrifugation and lysed by bead beating in buffer S6 (50 mM Tris-HCl, pH 7.5; 150 mM NaCl; 5 mM EDTA; 10% glycerol; 0.2% NP-40; 1 mM DTT; 1 mM PMSF; cOmplete Mini EDTA-free protease inhibitors [Roche]; 0.1 mM sodium orthovanadate; 20 μM sodium glycerophosphate; 20 μM para-nitrophenylphosphate; 20 μM sodium fluoride). Lysates were resolved by SDS-PAGE, transferred onto PVDF membranes, and probed for phosphorylated Rps6 using Ribosomal Protein S6/RPS6 antibody (Bio-Techne Canada). Beta-actin was used as a loading control (monoclonal antibody AC-15, Thermo Fisher). Secondary detection was performed using a goat anti-human IgG Fc secondary antibody (Invitrogen), and blots were imaged with a ChemiDoc Imaging System (Bio-Rad).

### Determination of intracellular Pi content

Free and total Pi levels were determined using a colorimetric molybdate assay, as previously described (46). Pi concentration was calculated by interpolation from a KH_2_PO_4_ standard curve and normalized to the total protein, determined using a Bradford assay kit (Bio-Rad).

### RNA-seq analysis

Overnight cultures of the *ino80* mutant and WT strains were diluted to an OD_600_ of 0.1 in 100 ml of fresh SC medium and grown at 30°C to logarithmic phase (OD_600_ of 0.6) under normoxia. Cells were then harvested by centrifugation, washed once with PBS, and resuspended in 400 μl of SC medium. Cultures were incubated at 30°C with shaking at 220 rpm for 15 min in 6-well plates under normoxic (21% O_2_) or hypoxic (5% O_2_) conditions, using the Cytation 5 plate reader equipped with O_2_ and CO_2_ gas controllers. Cells were then centrifuged for 2 min at 3,500 rpm and the pellets were rapidly frozen and stored at −80°C. Total RNA was purified using glass bead lysis in a Biospec Mini-beadbeater 24 and the RNeasy purification kit (Qiagen), as previously described (58). Expression analysis by RNA-seq was performed as previously described (59). cDNA libraries were prepared using the NEBNext Ultra II RNA Library Prep Kit, and sequencing was performed using an Illumina NovaSeq 6000 system at the Genome Quebec facility (Centre d’expertise et de services, Montreal). Gene ontology (GO) analysis was performed using FungiDB GO Enrichment Analysis (60).

### ChEC-seq analysis

ChEC-seq (Chromatin Endogenous Cleavage sequencing) analysis was performed as previously described by Tebbji *et al.* (61). *INO80* was MNase-tagged using a PCR cassette generated from the pFA-MNase-ARG4 plasmid. *C. albicans* MNase-tagged and free-MNase control strains were grown overnight in SC medium at 30°C. Cultures were then diluted to an initial OD_600_ of 0.1 in 50 ml of fresh SC medium and incubated at 30°C until reaching an OD_600_ of 0.5. 6-well plates were used to grow cells under either normoxic or hypoxic conditions for 15 min in the Cytation 5 plate reader. Cells were pelleted (3,000 × g for 5 min) and washed three times with 1 ml of buffer A (15 mM Tris-HCl [pH 7.5], 80 mM KCl, 0.1 mM EGTA, 0.2 mM spermine, 0.5 mM spermidine, protease inhibitor cocktail, and 1 mM PMSF). Cell pellets were resuspended in 800 μl buffer A containing 0.1% digitonin and permeabilized for 10□ min at 30°C with shaking. MNase activity was induced by adding CaCl_2_ to a final concentration of 5 mM and incubating samples at 30°C for 5 min. The reaction was quenched with 50 μl of 250 mM EGTA. DNA was purified using the MasterPure Yeast DNA Purification Kit and resuspended in 50µl of 10 mM Tris-HCl buffer (pH 8.0). RNA was digested with 10 μg of RNase A at 37°C for 20 min. Library preparation, sequencing, and peak calling were performed as described previously (61).

### Micrococcal nuclease digestion and nucleosome mapping

Chromatin preparation for micrococcal nuclease (MNase) digestion was performed as described previously (62). The contribution of Tor1 and Ino80 to chromatin accessibility was tested using *tor1* and *ino80* GRACE mutants under either repressible (20 µg/ml doxycycline) or non-repressible conditions as in the ChEC-seq experiment. *C. albicans* cells were crosslinked, and spheroplasts were subsequently treated with 6 U of MNase (Worthington Biochemical) for 30 min at 37°C. Reactions were terminated by adding 50 mM EDTA and 50 mM EGTA, followed by RNA removal using RNase A (0.25 mg/ml; Life Technologies). Samples were then de-crosslinked by overnight incubation at 60°C in the presence of 0.8% SDS and 1 mg/ml proteinase K (Invitrogen). DNA was purified using phenol-chloroform-isoamyl and ethanol precipitation. Sequencing libraries were prepared using the KAPA HyperPrep kit (Roche) and sequenced on an Illumina NovaSeq 6000 sequencing system at the Genome Quebec facility. Nucleosome occupancy and accessibility were determined using the nucMACC pipeline (63). Due to limitations in annotation, the ATG start codon was used as a proxy for the transcription start site (TSS).

### HT-29 damage assay

HT-29 enterocyte damage was assessed as previously described (23). HT-29 cells (ATCC HTB-38) were seeded as monolayers at 2 × 10^4^ cells per well in 96-well plates containing McCoy’s 5A medium supplemented with 10% fetal bovine serum and incubated at 37°C with 5% CO_2_for 24 h. Overnight cultures of *C. albicans* strains were pelleted, resuspended in cell culture medium, and added to HT-29 monolayers at a host-to-yeast multiplicity of infection (MOI) of 1:2. Co-incubations were performed for 24 hours at 37°C with 5% CO_2_ under normoxic or hypoxic conditions using a Heracell VIOS Tri-Gas incubator. After incubation, 100 μl of supernatant was collected from each well, and enterocyte damage was quantified by measuring lactate dehydrogenase (LDH) release using the Cytotoxicity Detection Kit PLUS (Roche).

### Galleria virulence assay

Last-instar *Galleria mellonella* larvae (180 ± 10 mg; Elevages Lisard, Saint-Bruno-de-Montarville, QC, Canada) were used for systemic candidiasis infection assays. Overnight cultures of *C. albicans* grown in SC medium were washed twice and diluted in PBS to 2.5 × 10^5^ cells per 20 μl. PBS alone was used as a negative control. Larvae were injected with 20 μl of the inoculum into the last left pro-leg and incubated at 37°C. Mortality was monitored daily for six days, with death defined as lack of response to touch and inability to right themselves. Survival data were analyzed using Kaplan-Meier curves and compared with the log-rank test (GraphPad Prism 10).

### Biofilm assay

Biofilm formation was assessed using XTT reduction assays (64). Overnight *C. albicans* cultures were washed three times with PBS and resuspended in RPMI 1640 supplemented with L-glutamine (0.3 g/l) to an OD_600_ of 0.1. Cells were allowed to adhere to flat-bottom 96-well polystyrene plates for three hours at 37°C with gentle rocking. Non-adherent cells were removed by three PBS washes, and fresh RPMI medium was added. Plates were incubated for 24 hours at 37°C under agitation (220 rpm) to allow biofilm formation. Wells were then washed three times with PBS, and 100 μl of XTT-menadione solution (0.5 mg/ml XTT in PBS with 1 mM menadione in acetone) was added. Following a three-hour incubation in the dark at 37°C, 80 μl of supernatant was collected for colorimetric measurement at OD_495_ to quantify biofilm metabolic activity. At least four biological replicates were performed.

### Statistical analysis

All phenotyping experiments were performed with at least three biological replicates. Gene expression and chromatin-profiling assays were conducted with two biological replicates. Statistical analyses and graph generation were carried out using GraphPad Prism (version 10.6.1).

Statistical difference between two sets of data with a non-parametric distribution was assessed using one-way ANOVA (Tukey’s multiple comparison test). The following p-values were considered: *p < 0.05; **p < 0.01; ***p < 0.001; ****p < 0.0001.

## Data availability

All RNA-seq, MNase-seq and ChEC-seq data are available in the GEO database (https://www.ncbi.nlm.nih.gov/geo/) under the accession number GSE314174.

## Supplementary data

**Supplementary Figure S1.** Validation of hypoxia-screen hits. Mutant strains were grown for 24 hours under normoxic and hypoxic conditions, and OD_600_ measurements were recorded every 10 min. Data represent the mean of three independent biological replicates.

**Supplementary Figure S2.** Relative hypoxic growth of the *pho4* (**A**) and *pho87* (**B**) mutants in media containing no, low, or high phosphate (Pi), normalized to WT growth under the same conditions.

**Supplementary Table S1.** List of *C. albicans* GRACE mutants whose growth is exacerbated or alleviated by hypoxia.

**Supplementary Table S2.** Biological functions of genes required for hypoxic growth.

**Supplementary Table S3.** RNA-seq analysis of the *ino80* mutant under hypoxia. Lists of raw RNA-seq data and differentially expressed transcripts in the *ino80* mutant compared to the WT strain under hypoxic conditions. Differential expression was determined using a false discovery rate (FDR) of 1% and a ±1-Log_2_ fold change cutoff.

**Supplementary Table S4.** List of Ino80 occupied loci identified by ChEC-seq under hypoxic conditions.

**Supplementary Table S5**. *C. albicans* hypoxia-modulated transcripts. Transcripts were identified by comparing the transcriptome of WT cells grown under hypoxia to that of cells grown under normoxia, using an FDR of 1% and a ±0.5-Log_2_ fold-change cutoff.

**Supplementary Table S6.** Primer and strain lists.

