## Supplementary figures and images for "Crosstalk between the Ino80 complex and TOR signaling drives fungal adaptation to hypoxia through chromatin remodeling"

### Supplementary Figure S1

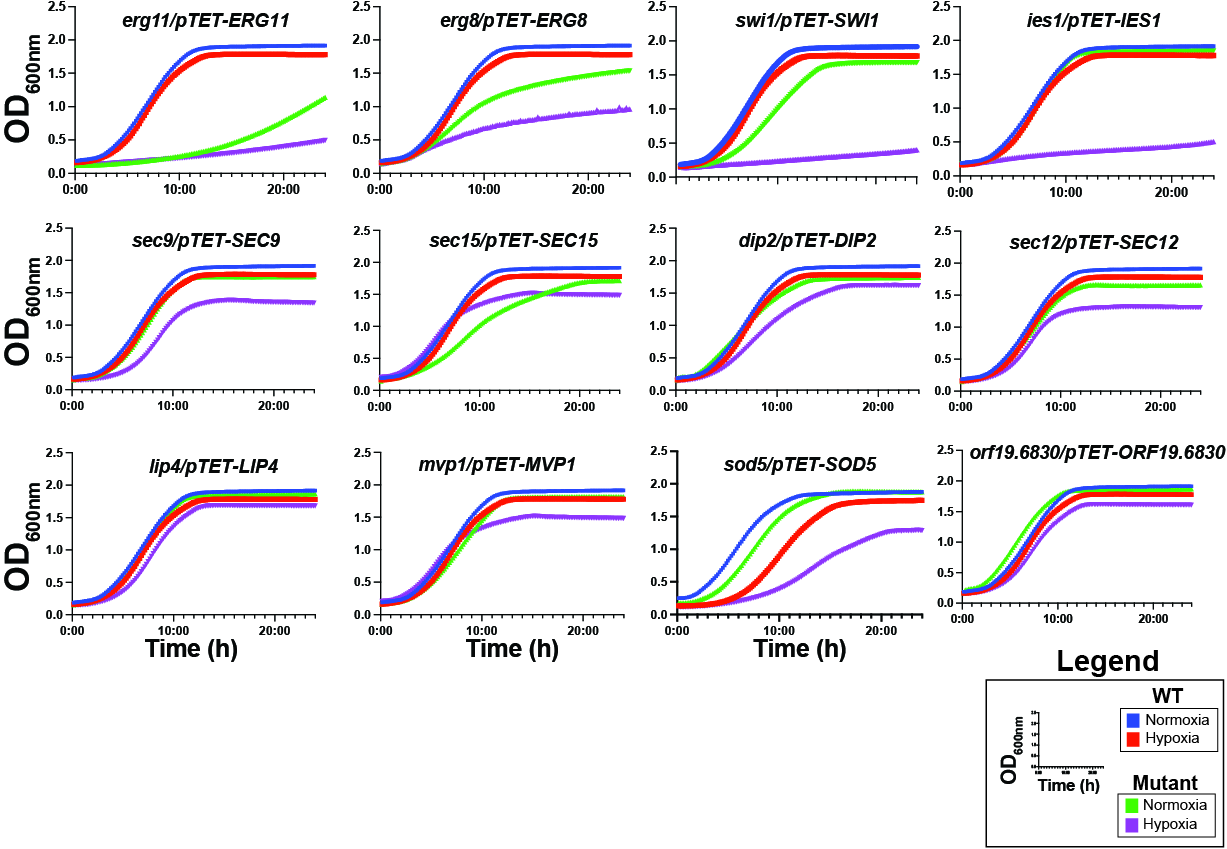

### Supplementary Figure S2

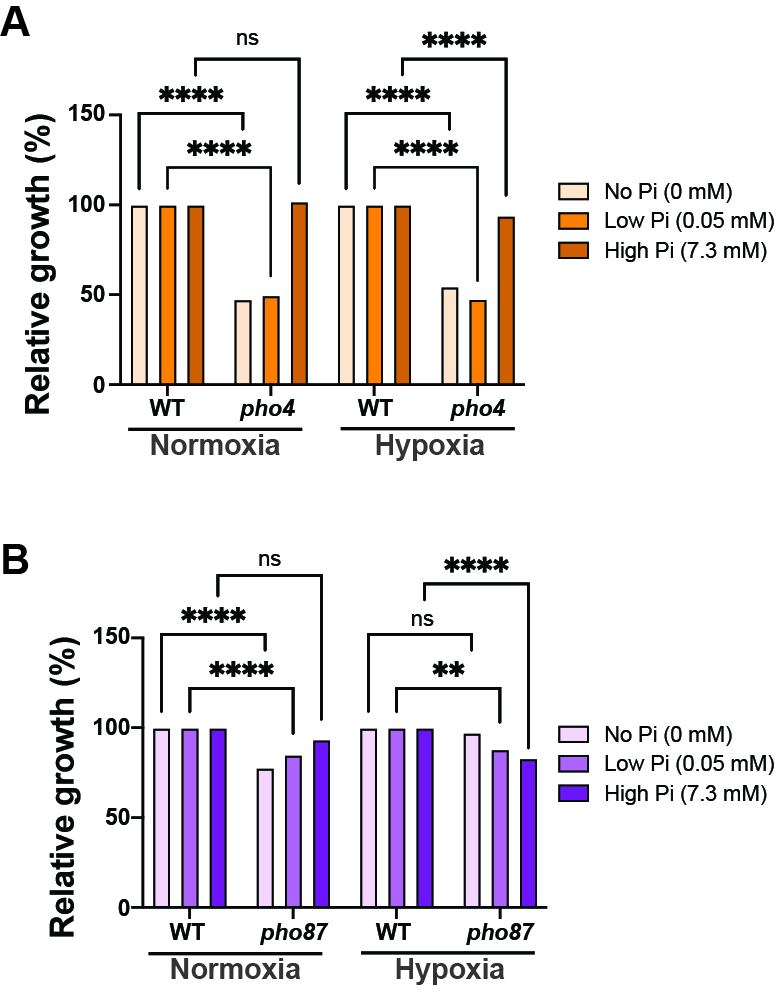
